# Reconstruction of the Cortical Maps of the Tasmanian Tiger and Comparison to the Tasmanian Devil

**DOI:** 10.1101/083592

**Authors:** Gregory S. Berns, Ken W.S. Ashwell

## Abstract

The last known Tasmanian tiger (*Thylacinus cynocephalus*) – aka the thylacine – died in 1936. Because its natural behavior was never documented, we are left to infer aspects of its behavior from museum specimens and unreliable historical recollections. Recent advances in brain imaging have made it possible to scan postmortem specimens of a wide range of animals, even more than a decade old. Any thylacine brain, however, would be more than 100 years old. Here, we show that it is possible to reconstruct white matter tracts in two thylacine brains. For functional interpretation, we compare to the white matter reconstructions of the brains of two Tasmanian devils (*Sarcophilus harrisii*). We reconstructed the cortical projection zones of the basal ganglia and major thalamic nuclei. The basal ganglia reconstruction showed a more modularized pattern in the cortex of the thylacine, while the devil cortex was dominated by the putamen. Similarly, the thalamic projections had a more orderly topography in the thylacine than the devil. These results are consistent with theories of brain evolution suggesting that larger brains are more modularized. Functionally, the thylacine’s brain may have had relatively more cortex devoted to planning and decision-making, which would be consistent with a predatory ecological niche versus the scavenging niche of the devil.

**SIGNIFICANCE:** The Tasmanian tiger – aka the thylacine – was a carnivorous marsupial and apex predator in Tasmania until the last one died in 1936. Very little is known about its natural behavior. Only four intact brain specimens are known to have survived. We used diffusion-weighted MRI to reconstruct the white matter pathways in two of these specimens, and for comparison, the brains of two Tasmanian devils. By comparing the maps of these projections in the cortex, we show that the thylacine’s brain is consistent with a more complex predatory strategy than the scavenging strategy of the Tasmanian devil.

## INTRODUCTION

The Tasmanian tiger (*Thylacinus cynocephalus*) – aka the thylacine – was, perhaps, the most iconic animal of Tasmania. A carnivorous marsupial, the thylacine was the apex predator in Tasmania until the last known animal died in the Hobart Zoo in 1936. The thylacine’s demise can be directly attributed to the bounty scheme in place from 1830-1914 that resulted in the killing of several thousand animals and indirectly to the loss of its habitat from farming activity (1). Although several animals had been kept in captivity in the early 1900s, no systematic investigation of the thylacine’s behavior was ever documented. The only records of behavior in their natural habitat are stories passed on by farmers, hunters, and trappers (2). Thus, in one of the great lost opportunities, very little is known about thylacine behavior (3).

Even without naturalistic data, it is still possible to reconstruct aspects of thylacine behavior from artifacts. The geometry of its elbow joint suggested that it hunted more by ambush than pursuit (4). Analysis of tooth morphology suggested the thylacine was a “pounce-pursuit” predator that killed prey in the 1-5 kg range (5). Analysis of skull mechanics have come to different conclusions about the size of the thylacine’s prey, with one analysis suggesting that the thylacine preyed on animals smaller than its size (6) while another suggested the opposite (7). This is in contrast to the scavenging strategy of the extant carnivorous marsupial, the Tasmanian devil (*Sarcophilus harrisii*).

To further understand thylacine behavior and to place the thylacine in its evolutionary context, we can look to brain morphology (8). Endocasts have suggested a more highly gyrified cortex than the Tasmanian devil, which is consistent with a greater encephalization quotient of the thylacine (0.45) than the devil (0.36) (9). A larger, more gyrified, brain might simply reflect the larger body size of the thylacine, or it might reflect a more sophisticated cognitive architecture, perhaps related to its predatory ecology.

To answer these questions, we used MRI and diffusion-weighted imaging (DWI) to reconstruct the architecture and white-matter pathways of an intact thylacine brain. Similar reconstructions in cetacean brains have shown this is possible in specimens more than a decade old (10). By comparing the thylacine brain structure to the Tasmanian devil, we can then infer structural-functional relationships between brain and behavior. Only four thylacine brains are documented to have survived intact (11). Here, we report the results from two of them, with comparison to the brains of two Tasmanian devils.

## MATERIALS AND METHODS

### Specimens

The brains of two thylacines and two Tasmanian devils were used (Fig. 1). Thylacine 1 was provided on loan by the Smithsonian. The history of the animal was well-known. It had been a male, one of two siblings living at the National Zoological Park until his death on Jan. 11, 1905. The brain was extracted by Ales Hrdlicka. The body weight was recorded as 14.97 kg, and the brain weighed 43 g. It was initially preserved in a solution of alum and 5% formalin. At some point, this was changed to the current solution of 1 part 37% formalin : 7 parts water : 12 parts 95% ethanol. Immediately prior to scanning, the brain weighed 16.8 g. Relatively little is known about Thylacine 2, which was provided on loan by the Australian Museum in Sydney. It weighed 30.6 g, was missing the olfactory bulbs, and had a large cut in the left dorsal cortex (Fig. 1B).

**Fig. 1.**
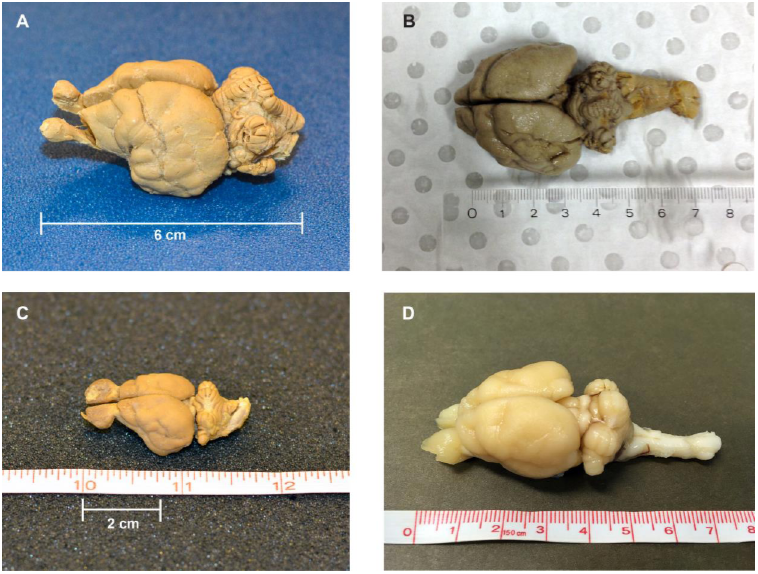
Specimens. A) Thylacine 1 (courtesy Smithsonian); B) Thylacine 2 (courtesy Australian Museum); C) Devil 1 (courtesy Smithsonian); D) Devil 2 (courtesy Save the Tasmanian Devil Program).

Devil 1 was also provided by the Smithsonian and was preserved using similar methods around the same time as Thylacine 1. Immediately prior to scanning, it weighed 4.6 g. Devil 2 was provided by the Save the Tasmanian Devil Program (STDP), a division of the Tasmanian Department of Primary Industries, Parks, Water and Environment (DPIPWE). The devil had been euthanized for age and health reasons. The extracted brain was preserved in formalin and ethanol and weighed 15 g immediately prior to scanning. Structural scans revealed a small cut near the right thalamus and brainstem.

For scanning, all specimens except Thylacine 2 were submerged in Fluorinert FC-3283 (3M). This minimized field distortions due to air/brain boundaries. Devil 1 was presoaked in phosphate buffered saline for 3 days to increase signal. Because of time constraints, Thylacine 2 was bagged and scanned.

### Imaging

Except for Thylacine 2, imaging was performed on a 3 T Siemens Trio with standard gradients and a 32- channel head receive coil. DTI was acquired with a diffusion-weighted steady-state free precession (DW-SSFP) sequence (12). Although DW-SSFP is more motion-sensitive than the usual diffusion-weighted spin-echo sequences used for *in vivo* DTI, DW-SSFP is more SNR efficient for tissues with short T2, which is critical for postmortem imaging. Because specimens varied in size and preservation quality, each required a different scanning protocol to optimize SNR.

For Thylacine 1, we acquired 12 sets of DW-SSFP images weighted along 52 directions (voxel size = 1.1 mm isotropic, TR = 26 ms, TE = 20 ms, flip angle = 37°, bandwidth = 159 Hz/pixel, q = 226 cm^−1^, G_max_ = 38.0 mT/m, gradient duration = 14.0 ms). Twenty images were acquired with these same parameters except with q = 10 cm^−1^ applied in one direction only (these serve as a signal reference similar to b=0 scans in conventional spin-echo acquisitions). Proper modeling of the DW-SSFP signal requires knowledge of T1 and T2 values, which are drastically altered in postmortem compared to *in vivo* tissue. These values were calculated based on a series of T1-weighted images (TIR sequence with TR = 1000 ms, TE = 12 ms, and TI = 30, 120, 240, 900 ms) and T2-weighted images (TSE sequence with TR = 1000 ms, TE = 15, 31, 46 ms). Structural images were acquired using a 3D balanced SSFP sequence (TR = 8.5 ms, TE = 4.25 ms, flip angle = 60°). Balanced SSFP images were acquired in pairs with the RF phase incrementing 0° and 180°, which were averaged later to reduce banding artifacts (12, 13). This yielded structural images with 0.33 x 0.33 x 0.33 mm resolution. It took approximately 6 hours to acquire all scans.

For Devil 1, the same protocol was used with the following change in parameters: 10 sets of DW-SSFP images (51 directions, voxel size = 1.5 mm isotropic, TR = 22 ms, TE = 17 ms, flip angle = 50°, bandwidth = 373 Hz/pixel, q = 205 cm^−1^, gradient duration = 12.7 ms). For Devil 2: 4 sets of DW-SSFP images (52 directions, voxel size = 1 mm isotropic, TR = 31 ms, TE = 24 ms, flip angle = 29°, bandwidth = 159 Hz/pixel, q = 255 cm^−1^, gradient duration = 15.8 ms). For Devil 2, we acquired a high resolution structural image using a turbo-spin echo sequence which provided better contrast (2 averages, voxel size = 0.33 mm isotropic, TR = 1000 ms, TE = 50 ms).

Thylacine 2 was scanned on a Bruker 9.4 T BioSpec. For DTI, one set of spin-echo diffusion-weighted volumes were acquired with the following parameters: 45 slices oriented in the sagittal plane with in-plane resolution of 1 mm, slice thickness = 0.78 mm, 30 gradient directions with b_eff_ = 2050 s/mm^2^, and 5 volumes with b_eff_ = 5 s/mm^2^ to serve as a b0 reference. For anatomical reference, we acquired a T2-weighted spin-echo image with an in-plane resolution of 0.1 mm and between-plane resolution of 0.2 mm (2 acquisitions).

Thylacine 1 and Devils 1 & 2 were processed with FSL tools modified to account for the DW-SSFP signal model and which incorporated tissue T1 and T2. All diffusion images were registered to one q=10 cm^−1^ reference image, and the references were averaged together to create a mean reference. Transformations between the structural image and mean DTI were used to map regions-of-interest (ROI) for tractography. To fit a diffusion tensor model at each voxel, we fitted an extension of the model proposed by Buxton (14) to incorporate Gaussian (DTI) anisotropic diffusion (15). The tensor parameters (3 eigenvalues, 3 orientations) were estimated using the Metropolis Hastings algorithm with a positivity constraint of the tensor eigenvalues. This yielded estimates for fractional anisotropy, mean diffusivity, and three eigenvectors representing the principal directions of the diffusion tensor (15). For tractography, we used a similarly modified version of BEDPOSTX that incorporated the new signal model (13, 14, 16) with 2 crossing fibres per voxel but otherwise default options (17). Thylacine 2 was processed with the unmodified FSL tools. Relaxation parameters and b_eff_ are shown in Table 1. Because the orientation of the specimens within the magnet varied and was different than would be standard for a human brain, this required reordering and inversion of some directions in the vector file. Vector directions were confirmed by verifying that major tracts (e.g. anterior commissure, fornix, brainstem) were oriented correctly.

**Table 1.** Relaxation parameters and b_eff_.

| Specimen | T1 (ms) | T2 (ms) | $b_{\text{eff}}$<br>(s/mm<sup>2</sup>) |
| --- | --- | --- | --- |
| Thylacine 1 | 150 | 30 | 1500 |
| Thylacine 2 | N/A | 30 | 2050 |
| Devil 1 | 120 | 30 | 950 |
| Devil 2 | 550 | 60 | 3000 |

### Tractography

Because little is known about the cortical maps of the thylacine, we performed probabilistic tractography to determine the cortical fields of the basal ganglia and major thalamic nuclei. In each specimen, the basal ganglia were divided into 4 regions of interest (ROIs): 1) dorsal caudate; 2) mid caudate; 3) ventral caudate (including nucleus accumbens); and 4) putamen (Fig. 2). ROIs were drawn on the structural images. The cortex of each hemisphere was masked separately and served as the seed ROI for *probtrackx2*, using the subcortical ROIs as targets. The procedure was done for the thalamus using 5 ROIs (Fig. 3): 1) dorsolateral geniculate (DLG); 2) medial geniculate (MGN); 3) lateral posterior thalamic nucleus (LP); 4) ventral anterior and ventrolateral nuclei (VAVL); and 5) ventral posterior lateral and medial nuclei (VPLM). Thalamic nuclei can be challenging to visualize and were based on the known anatomy of the brains of other carnivorous marsupials (18).

**Fig. 2.**
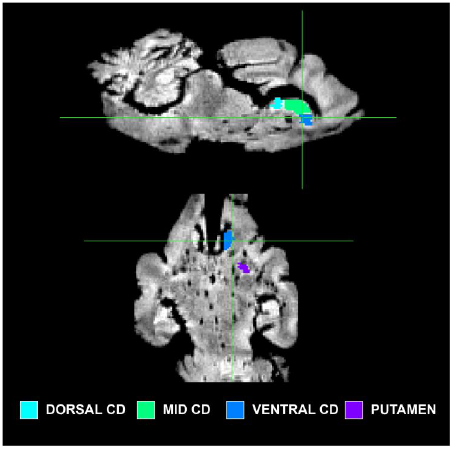
Subdivisions of the basal ganglia in Thylacine 1.

**Fig. 3.**
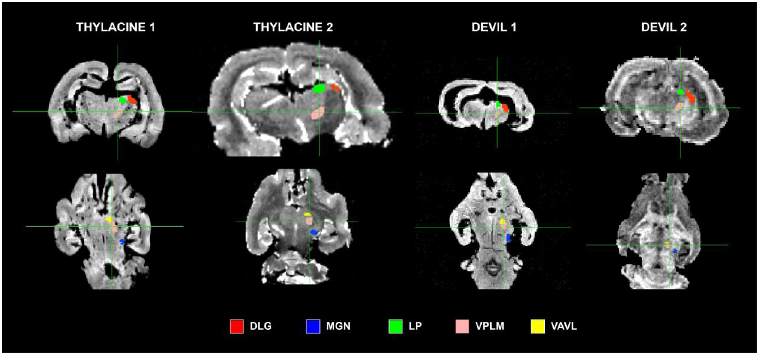
Location of thalamic ROIs.

For every voxel in these seed masks, a series of “streamlines” were calculated that formed a representation of the likely tract structure by incorporating the underlying uncertainty in the preferred diffusion of water, which occurs along the predominate orientation of fibers passing through a voxel (12). By proceeding from voxel to voxel, one can trace specific fiber tracts. We then used *find_the_biggest* to create the cortical map in which each voxel was mapped to the ROI with the greatest number of streamlines going through it. This was done for both the basal ganglia and thalamic nuclei. 3D renderings of the cortical maps were performed with ITK-SNAP.

For 3D visualization of the white matter tracts, we used deterministic tractography as implemented in DSI Studio (19). This algorithm estimates the orientation distribution functions (ODFs) by first filtering out noisy fibres based on a quantitative anisotropy (QA) metric. We used this approach primarily for visualization and as a complement to the probabilistic methods implemented in FSL. For whole-brain rendering, we used the following parameters: QA threshold = 0.07, angular threshold = 55°, step size = 0.55 mm, minimum length = 10 mm. 3D images of tracks were rendered at 1° increments of rotation of the brain and assembled into a movie. The ODF estimated in DSI Studio is theoretically only a valid fiber ODF in the case of standard pulsed-gradient spin-echo diffusion experiments, and may not represent the true ODF for SSFP diffusion experiments. However, for the purposes of tractography, we only use the ODF peaks, not the full ODF shape, and therefore, this model-free ODF approach remains a valid approach even for SSFP diffusion, especially for visualization.

## RESULTS

Qualitatively, the 3D visualizations of the devil and the thylacine showed the effectiveness of DTI, even in specimens over 100 years old (Fig. 4). Devil 2 (Movie 1), which was the fresh specimen, displayed tract reconstructions broadly consistent with marsupial brain anatomy (18) and served as the best reference for the other specimens. A broad swath of fibers going left-right (*red*) constituted the anterior commissure and can be seen sweeping dorsally to the cortex. These fibers are intermingled with the internal capsule, whose fibers can be distinguished by their caudal trajectory (*green*). Tracts to the olfactory bulbs are clearly identified. The fibers of the fornix and fimbria of the hippocampus can be seen as a ‘V’ emanating dorsal to the anterior commissure and arching dorsolaterally. Caudally, the left-right fibers of the pons are easily identified, as are the tracts of the cerebellar peduncles.

**Fig. 4.**
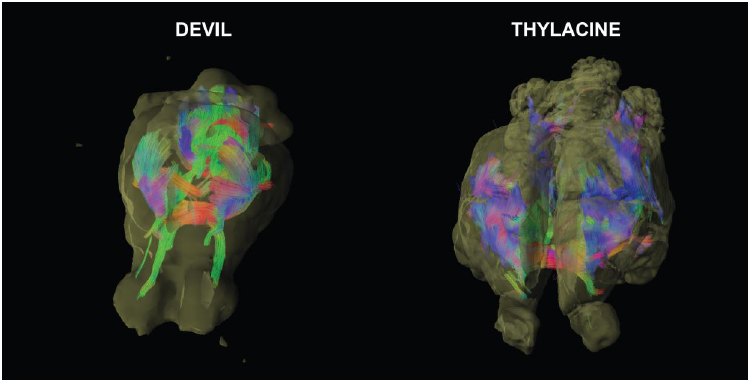
3D reconstructions of the white matter tracts of the devil (Movie 1) and thylacine (Movie 2).

Thylacine 1 (Movie 2), although 100 years old, still showed identifiable tracts. As in the devil, the anterior commissure dominated the interhemispheric connections, but because the cortex is larger in the thylacine, the upsweep of fibers appears more prominent. The internal capsule fibers can also be seen, although the cortical target is somewhat more rostral than in the devil. The fornix and fimbria, although discernable, were not as prominent as in the devil. In general, the long tracts appeared more segmented in the thylacine – likely due to its age.

The basal ganglia pathways showed broad similarities in all of the specimens (Fig. 5). The caudate ROIs – dorsal, mid, and ventral – maintained this organization along a medial swath of cortex, with some lateral spread caudally. The putamen pathways were located just lateral to the caudate zones and represented the likely location of motor regions. Because the left cortex of Thylacine 2 was damaged, pathways were displaced laterally but preserved the same topography.

**Fig. 5.**
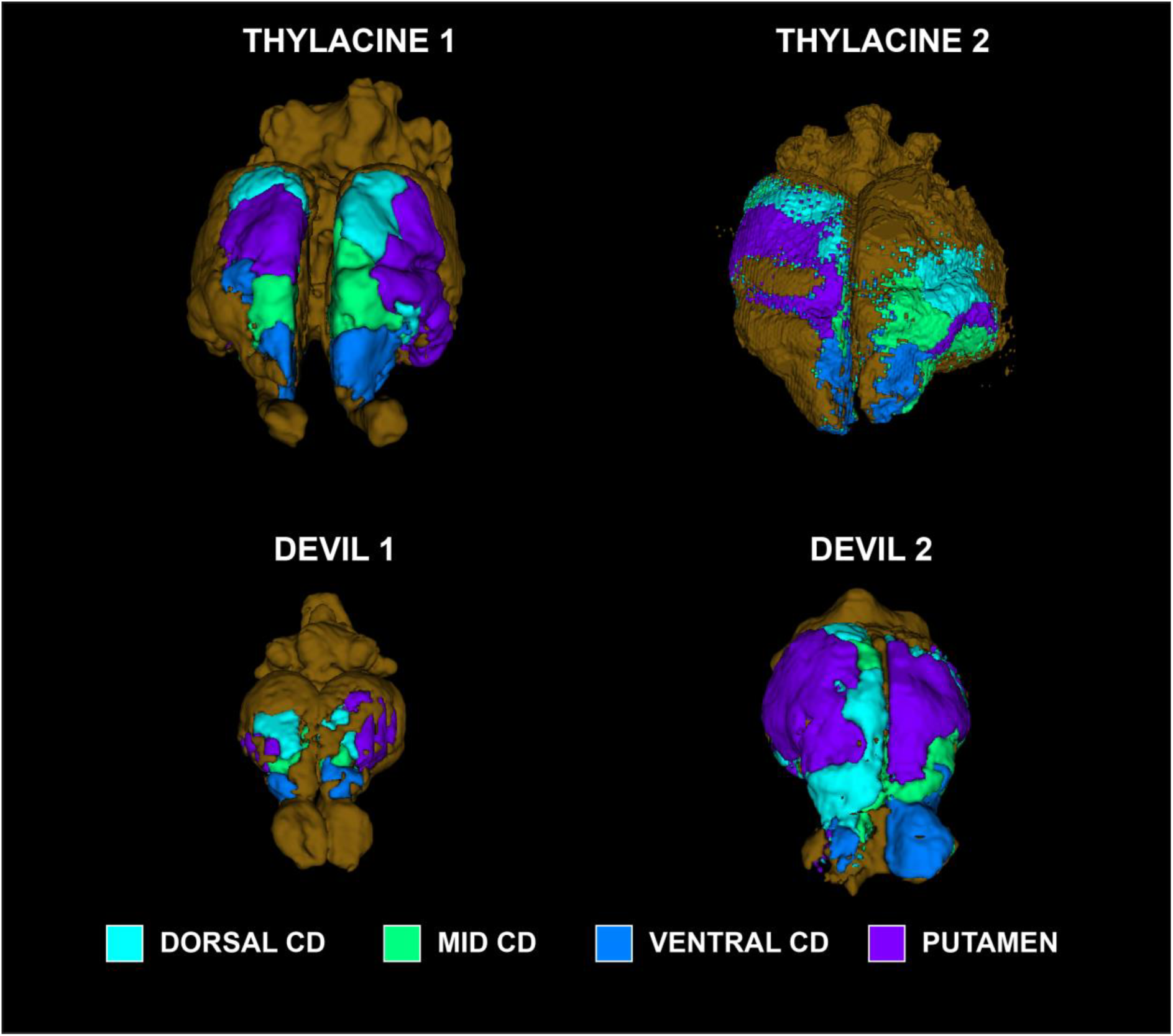
Cortical maps of basal ganglia projections.

The thalamic maps had similar general layouts, although each of the specimens differed in the maximal cortical locations of the thalamic nuclei (Fig. 6). Thylacine 2 had damage to the left dorsal cortex and right thalamus, so the right cortical map was mostly nonexistent and the left map was displaced laterally as with the BG. The DLG projection was located dorsally in all other specimens. The MGN projection extended from the caudal cortex to the ventral surface to varying degrees in all specimens. In Devil 2, the VPLM seed dominated much of the cortical map. The LP occupied only a small portion of cortex and in variable locations. However, both VAVL and VPLM mapped to consistent locations with the VAVL zone located rostrally, with the VPLM zone just caudal and lateral to VAVL (except in Devil 2). These zones would correspond roughly to motor (VAVL) and somatosensory regions (VPLM).

**Fig. 6.**
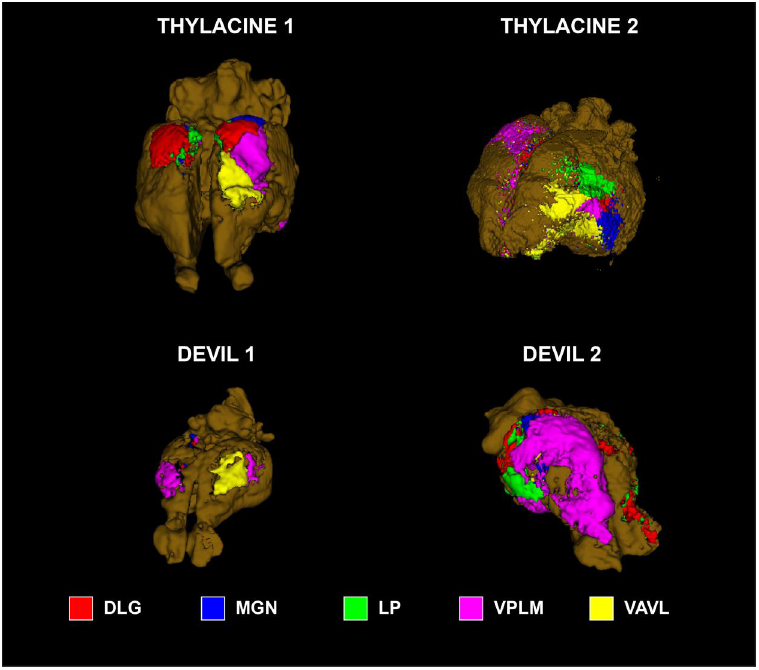
Cortical maps of thalamic nuclei. DLG=dorsolateral geniculate nucleus, MGN=medial geniculate nucleus, LP=lateral posterior nucleus, VPLM=ventrolateral/medial posterior nucleus, VAVL= ventral anterior and ventrolateral nuclei.

## DISCUSSION

The results presented here show, for the first time, the reconstruction of white matter pathways in the brain of the thylacine – aka Tasmanian tiger. The last known thylacine died in 1936, and only four preserved brains are known to exist. Here, we present the results from two of them. Performing DTI in such specimens is challenging because of their age, with each approximately 100 years old. Despite the age, we were able to reconstruct major pathways between the cortex and the basal ganglia and the thalamus. In comparison to the thylacine’s closest living relative – the Tasmanian devil – we find broad similarities with some differences that inform theories of brain evolution.

The age of the specimens places limitations on the fidelity of information we can extract. 100 years in preservative takes a toll. This was particularly striking in the Smithsonian thylacine. Because of excellent record keeping, we know the original weight of the brain was 43 g. Prior to scanning it was 16 g. Although we don’t know the history of storage and preservative changes, we can estimate that the specimen had shrunk at the rate of 1%/year. At such a rate, after 110 years, the specimen would weigh 33% of the original weight. This raises the question of how such shrinkage affects the white matter topology. If it shrunk uniformly, then the topology would be preserved. To examine the effects of age, we procured the brain of a Tasmanian devil from the Smithsonian’s collection that was preserved around the same time as the thylacine. We were also able to obtain the brain of a devil that had been recently euthanized. By comparing the white matter reconstructions of these two specimens, we can get a qualitative idea of what happens over a century in preservative.

Like the thylacine brain, the Smithsonian devil had shrunk substantially. Although we don’t know its original weight, it was roughly 1/3 the weight of the fresh specimen, suggesting a comparable degree of shrinkage as the thylacine. Comparing the two devil specimens, we see broad similarity in the cortical maps of the basal ganglia. Both specimens show the same topography of the dorsal, mid, and ventral caudate zones in a caudal to rostral direction over the dorsal surface of the cortex. The putamen zone is somewhat rostral and lateral. The fresh devil (Devil 2) had larger zones, which we assume is due to the higher fidelity signal. In addition to the shrinkage, the older specimens had a substantially shortened T1 and T2. With a T1=150 ms and T2=30 ms, the scan parameters had to be altered to achieve reasonable SNR. The TR was shortened, but this required a compromise in the strength of the diffusion gradient to stay within the power cycling limit of the scanner. Thus, the b_eff_ was somewhat less than optimal for some specimens.

Despite these limitations, the cortical maps were broadly consistent with both electrophysiological recordings and tract tracing studies in other marsupials and monotremes (20–22). We found that the motor system, as mapped by the cortical zone of the putamen, represented consistently on the lateral surface of the rostral half of the cortex. This motor zone was consistent with the cortical fields obtained from the ROI of thalamic nuclei VA/VL, even though the thalamic tracts were more susceptible to noise because of their small size. Similarly, the somatosensory region, as indicated by the VPL/VPM fields, was consistently located lateral and ventro-caudal to the motor regions in all of the specimens.

Despite the age of the specimens, the overall consistency of the cortical fields demonstrated that the white matter tracts were sufficiently intact to reconstruct major pathways. The consistency between Devil 1 (100 years old) and Devil 2 (1 year old) showed that even though the old specimens had shrunk with age, they must have shrunk uniformly enough to avoid changing the geometry. The exception was Thylacine 2, but that had gross damage extending from the left cortex to the right thalamus. The tract reconstructions for that specimen were noisier than the others. Because of time limitations with that specimen we were able to collect only 30 diffusion directions, which was less than the others. Even though Thylacine 2 was scanned at 9 T, this underscores the importance of acquiring as many directions as possible.

Comparing the thylacine to the devil, a few differences are apparent. The basal ganglia maps suggest the putamen projections occupy a relatively larger percentage of cortex. This is a qualitative observation. With only two specimens of each species, it is not possible to do meaningful statistical comparisons. Because these maps were created with a “winner-take-all” algorithm, they don’t capture the degree of overlap between the fields. In fact, there is substantial overlap, and the relative size of the putamen field may indicate greater overlap with sensory fields in the devil. This would be consistent with theories of brain evolution suggesting that cortical fields become more modularized as the cerebral cortex gets bigger (23). There is some hint of this in the thalamic maps as well. Devil 2, in particular, showed greater interdigitation of the thalamic fields in the cortex but was also dominated by the VPL/VPM fields on the right, which may have been affected by the cut in the thalamus on that side.

Overall, these findings show that it is possible to use DWI to reconstruct white matter pathways in brains that are over a century old. In the case of the thylacine, they inform structure-function relationships that would otherwise be impossible. The differences from the devil suggest a more modularized cortex, which may be due to its larger size. However, the relatively larger caudate zones in the cortex, particularly the mid and ventral portions, suggest a greater portion of cortex devoted to complex cognition especially related to action planning and, possibly, decision making. This would be consistent with the thylacine’s ecological niche, where hunting required more planning than the scavenging strategy of the devil.

## ACKNOWLEDGEMENTS

We are grateful to the Smithsonian Institution for loaning the thylacine and devil brains. In particular, Esther Langan and Darrin Lunde of the Mammal Division facilitated the loan process. Sandy Ingleby, at the Australian Museum in Sydney, made possible the scanning of their thylacine brain. Carolyn Hogg and the Save the Tasmanian Devil Program donated the brain of a devil. Stephen Sleightholme, maintainer of the International Thylacine Specimen Database, helped locate thylacine brains. Marco Gruwel assisted with the scanning at UNSW, and Peter Cook assisted with scanning at Emory University. Karla Miller and Saad Jbabdi at the FMRIB Centre, University of Oxford, made available the DWSSFP sequences and software modifications for reconstruction. Scan time was supported by Emory University and UNSW.

